# Within Thermal Scales: The Kinetic and Energetic Pull of Chemical Entropy

**DOI:** 10.1101/2023.09.20.558706

**Authors:** Josh E. Baker

## Abstract

Biological systems are fundamentally containers of thermally fluctuating atoms that through unknown mechanisms are structurally layered across many thermal scales from atoms to amino acids to primary, secondary, and tertiary structures to functional proteins to functional macromolecular assemblies and up. Understanding how the irreversible kinetics (i.e., the arrow of time) of biological systems emerge from the equilibrium kinetics of constituent structures defined on smaller thermal scales is central to describing biological function. Muscle’s irreversible power stroke – with its mechanochemistry defined on both the thermal scale of muscle and the thermal scale of myosin motors – provides a clear solution to this problem.

Individual myosin motors function as reversible force-generating switches induced by actin binding and gated by the release of inorganic phosphate, P_i_. As shown in a companion article, when *N* individual switches thermally scale up to an ensemble of *N* switches in muscle, the entropy of a binary system of switches is created. We have shown in muscle that a change in state of this binary system of switches entropically drives actin-myosin binding (the switch) and muscle’s irreversible power stroke, and that this simple two-state model accurately accounts for most key aspects of muscle contraction. Extending this observation beyond muscle, here I show that the chemical kinetics of an ensemble of *N* molecules differs fundamentally from a conventional chemical analysis of *N* individual molecules, describing irreversible chemical reactions as being pulled into the future by the a priori defined entropy of a binary system rather than being pushed forward by the physical occupancy of chemical states (e.g., mass action).

## I. INTRODUCTION

In 1824, studying the efficiency of heat engines, Carnot demonstrated that the mechanical state of a gas cylinder is not defined by the mechanical states of individual gas molecules in that cylinder [1]. In 1938, studying the efficiency of muscle contraction, A.V. Hill similarly demonstrated that the mechanical state of muscle is not defined by the mechanical states of individual molecules in muscle [2]. This extraordinary result has been largely ignored for the past 85 years because it is inconceivable to most molecular biologists that the function of a complex biological system is not defined classically by the function of proteins within that system [3–6]. Understanding how the top-down physics of Carnot and Hill converges with the bottom-up philosophy of molecular biologists is an open problem in biology and physics. Muscle – with mechanics and energetics well defined on two different thermal scales – provides a clear solution.

On the thermal scale of muscle, A.V. Hill showed experimentally that a system energy, *E*(*F*), in muscle is a linear function of muscle force, *F*, and is the energy available for muscle force generation and power output [2]. On the thermal scale of individual myosin motors, we have experimentally shown that myosin motors function as force-generating switches induced by actin binding and gated by inorganic phosphate release [7–11]. A.V. Hill demonstrated that the former is not classically defined by the latter, and we have shown that this is on account of the entropy that is created when *N* individual switches transform into an ensemble of *N* switches. The ensemble entropy created is that of a binary system of *N* switches well defined by statistical mechanics [12] and nonexistent in *N* individual switches (i.e., in *N* isolated single molecule mechanics experiments). While multiscale entropy approaches are widely used [13], they have not been explicitly described as thermal scaling.

In a companion article I show that thermal scaling physically occurs when we define the energetic state of an ensemble at which point the number of ways molecules within that ensemble can account for the ensemble state contributes to the ensemble state as entropy. At this point, the statistical occupancy of molecular states physically replaces the physical occupancy of molecular states, which is to say that thermal scaling is a quantum (on the scale of thermal energy), not a classical, phenomenon. In that article, I show that the ensemble of molecules is a resonant structure; that chemical transitions within the ensemble occur with tunneling through entangled barriers not classical diffusion over structural barriers; and that there is observational uncertainty in measuring the state of a molecule on the scale of the ensemble.

Here I develop the chemical energetics and kinetics of thermal scaling. I show that entropy created by thermal scaling defines the irreversible energetics and kinetics of the ensemble of molecules. Specifically, a change in the chemical state of the ensemble occurs with a change in ensemble entropy, which means chemical transitions are energetically driven by changes in ensemble entropy. I show that chemical kinetics is conventionally defined on the thermal scale of *N* molecules and that this differs fundamentally from the chemical kinetics defined on the thermal scale of an ensemble of *N* molecules. The former describes reversible chemical reactions being pushed into the future by the physical occupancy of chemical states defined in the present (i.e., mass action), whereas the latter describes irreversible chemical reactions being pulled into the future down an a priori defined entropic funnel. I show how with thermal scaling the former smoothly transforms into the latter and how the latter describes the arrow of time being pulled stochastically into a priori defined futures, where futures having more possibilities are kinetically and energetically more favorable.

Some may think this fantastical because it is at odds with the popular perspective that both scaling (e.g., entanglement and observational uncertainty) and entropy (Planck referred to it as a “spook”) are mysterious [14–17]. However, neither is mysterious; both thermal scaling and the usefulness of the entropy it creates are observed in muscle [8,9,11]. A.V. Hill’s 1938 observation disproved all classical mechanisms of biological function [2]. We have since demonstrated that the physical basis for Hill’s observation is the useful ensemble entropy physically created through thermal scaling, and that within the thermal scale of muscle an increase in this entropy with actin-myosin binding energetically drives the irreversible contraction of muscle [11] (the arrow of time). The implied two-state thermodynamic model accurately accounts for most key aspects of muscle contraction [9,11,18–20]. In short, thermal scaling and entropy are not mysterious. The mystery originates from a failure to understand that thermal scaling is a physical transformation. Once we account for this, entropy along with Schrodinger’s cat and Maxwell’s demon are no longer spooks, and with every heartbeat, every breath we take, and every step we make, entropy kinetically and energetically pulls us irreversibly into the future.

## II. MUSCLE: AN ENSEMBLE OF FORCE-GENERATING MOLECULAR SWITCHES

I use muscle – with energetics and kinetics well defined on two different thermal scales – to illustrate how the chemical kinetics of *N* individual molecules thermally scale up to the chemical kinetics of an ensemble of *N* molecules. On the thermal scales of individual myosin motors, a reversible myosin motor switch induced by actin binding and gated by the release of inorganic phosphate, P_i_ (state A to state B in Fig. 1A) generates force. On the thermal scale of muscle, a system energy, *E*(*F*), that varies linearly with muscle force, *F*, is the energy available for muscle force generation and power output (Fig. 1B). The former does not classically scale up to the latter because the latter physically contains the ensemble entropy of a binary system of switches and the former does not. In a companion article I describe in general how *N* molecules thermally scale up to an ensemble of *N* molecules. Here I describe the implications for chemical kinetics.

**Fig. 1.**
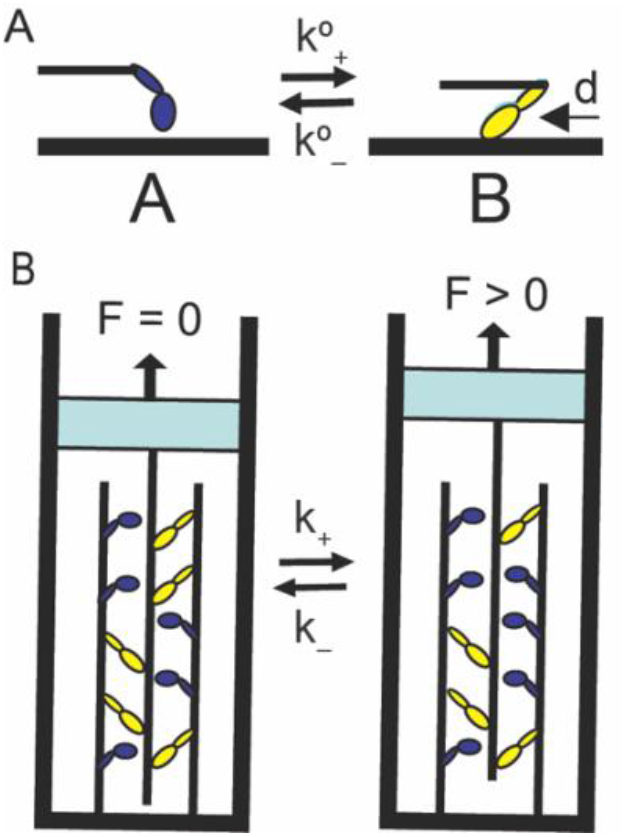
Entropic Force in a Binary Mechanical System. **(A)** A reversible molecular switch undergoes a discrete conformational change from chemical state A to B displacing compliant elements external to the switch a distance, *d*. The switch occurs with forward and reverse rates, *k*°_*+*_ and *k*°_*–*_. **(B)** An ensemble of molecular switches constitutes a binary mechanical system where, according to Le Chatelier’s principle, a force applied to the system reverses the force generating chemistry of the molecular switches.

## III. THE ENERGY LANDSCAPE OF AN INDIVIDUAL SWITCH

The change in free energy of a molecular switch with a chemical transition is defined by the energy landscape of that molecule (Fig. 2A), which a priori describes changes in its free energy along the molecule’s reaction coordinate{Formatting Citation} independent of the structural components that thermally scale up to create that molecule (see accompanying paper). The probability of a forward (A to B) relative to a reverse transition is *k*°_*+*_/*k*°_*-*_= exp(– ΔG°/k_B_T) [22], where *k*°_*+*_ and *k*°_*–*_ are forward and reverse rate constants and ΔG° is the standard (molecular) reaction free energy [23]. The standard reaction enthalpy, ΔH°, and entropy, ΔS°, contribute to the standard free energy as ΔG° = ΔH° – TΔS°, where ΔS° and ΔH° define the reaction entropy and enthalpy on the thermal scale of a molecule.

**Fig. 2.**
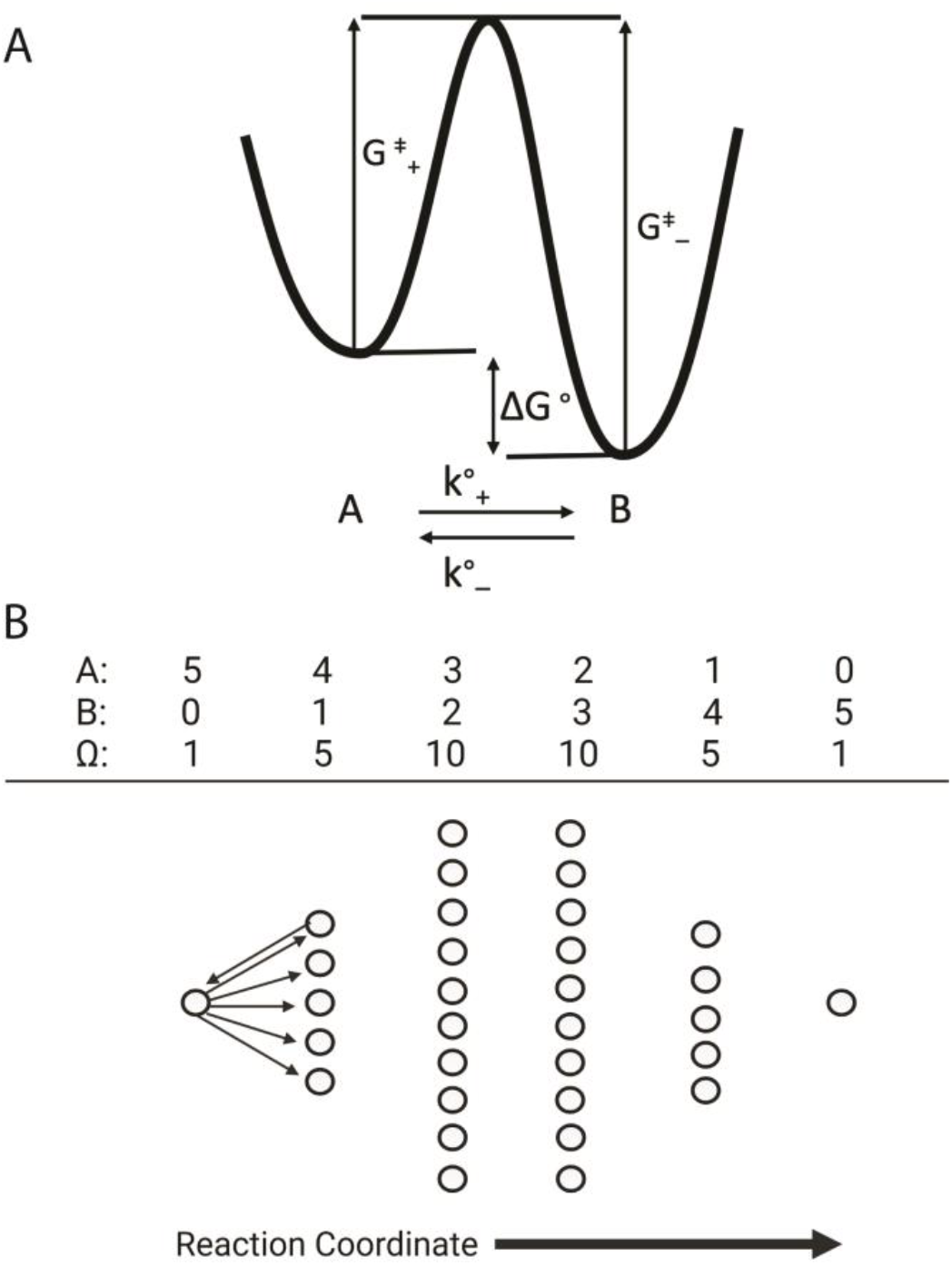
The molecular and system contributions to the reaction free energy. **(A)** The molecular contribution to the reaction free energy is described by a molecular energy landscape. The free energy difference between the two metastable states, A and B, is ΔG°. The rate, *k°*_*+*_, at which a molecule transitions from molecular state A to B varies exponentially with –G^‡^_+_, where G^‡^_+_ is the height of the activation barrier in the forward direction. The rate, *k°*_–,_ at which a molecule transitions from B to A varies exponentially with –G^‡^_–_, where G^‡^_–_ is the height of the activation energy barrier in the reverse direction. (**B**) The system contribution to the reaction free energy is entropic. For *N* = 5, there are 6 system states, {*N*_*A*_,*N*_*B*_}, along the reaction coordinate each with Ω_NA,NB_ = *N*!/(*N*_*A*_!*N*_*B*_!) microstates, illustrated with circles, and entropy k_B_·ln(Ω_NA,NB_). There are five micropathways (right arrows) by which a molecule in state {5,0} can transition to state {4,1}, and one micropathway (left arrow) by which a molecule in state {4,1} can transition to state {5,0}.

## IV. MOLECULAR AND ENSEMBLE FREE ENERGY

The state of *N* individual switches, *N*_*A*_ and *N*_*B*_, describes the physical occupancy of states A and B. The state of an ensemble of *N* switches,{*N*_*A*_,*N*_*B*_}, describes a probability density function centered at *N*_*A*_ = *N* – *N*_*B*_ and defined a priori for each state (see below). Here I define δη as a generic reaction coordinate that describes changes in either the ensemble state from {*N*_*A*_,*N*_*B*_} to {*N*_*A*_–δη,*N*_*B*_+δη} or molecular state from *N*_*A*_ and *N*_*B*_ to *N*_*A*_–δη and *N*_*B*_+δη.

Ignoring for simplicity mechanical (PV and non-PV work) contributions, the free energy, G, of any system in a given state is

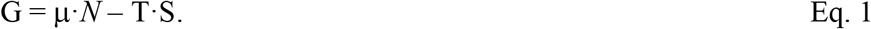

Equation 1 applies at every thermal scale from electron spins to amino acids to primary, secondary, tertiary, and quaternary protein structures to protein ensembles and assemblies and beyond, where the definition of the number, *N*, of components of the system, the chemical potential, μ, of those components, and the entropy, S, of an ensemble of those components physically changes across thermal scales. That is, as described in a companion article, G physically transforms across thermal scales. To better understand the mechanism for this physical transformation, here I consider how the free energy for *N* individual switches transforms into the free energy for an ensemble of *N* switches. I assume that P, V, T and N remain constant across this transition. The only variables in Eq. 1 are then μ, S, and G.

A change in system free energy, ΔG, with a change in state δη, is referred to as the reaction free energy

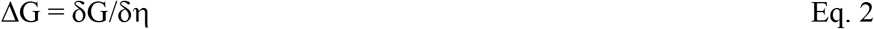

## V. A CHANGE IN CHEMICAL POTENTIAL, μ·N

When individual molecules, *N*_*A*_ and *N*_*B*_, are added to a system, the effect on free energy resembles that of a mixture of two gases, and μ·*N* = μ_NA_*N*_*A*_ + μ_NB_*N*_*B*_. When (*N*_*A*_ – 1) and (*N*_*B*_ + 1) individual molecules are added to a system, μ·*N* = μ_NA_(*N*_*A*_–1) + μ_NB_(*N*_*B*_+1). Thus, a change in state of one switch, δη = 1, results in a change in μ·*N* of ΔG° + k_B_T·ln(*N*_*B*_/*N*_*A*_), where ΔG° is the standard (molecular) free energy and *N*_*A*_ and *N*_*B*_ are the physical occupancies of states A and B commonly described as chemical activities. It is easy to see mathematically why a change in μ·*N* is not defined when *N*_*A*_ = 0 for forward reaction and when *N*_*B*_ = 0 for a reverse reaction; however, physically this does not make sense. Thermodynamics applies when *N*_*B*_ = *N* or *N*_*A*_ = *N* molecules are added to the system. The problem is that thermodynamic systems are defined by ensemble not molecular states.

As described in a companion article, an ensemble of *N* molecules cannot be divided into *N*_*A*_ and *N*_*B*_ molecules. When δη = *N*, the change in μ·*N* is the molecular free energy, ΔG°, summed over all *N* chemical steps. For δη = 1, the change in μ·*N* is then the molecular free energy, ΔG°, which is defined for both *N*_*A*_ = 0 and *N*_*B*_ = 0. In summary, for a single chemical step, δη = 1, of a system of *N* individual molecules

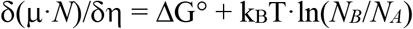

whereas for a single chemical step, δη = 1, of an ensemble of *N* molecules,

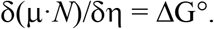

Thus, as described in a companion article, with thermal scaling from *N* individual molecules to an ensemble of *N* molecules, chemical activities and the physical occupancy of molecular states are physically lost from the system.

## VI. ENSEMBLE ENTROPY, ΔS: THE EQUIVALENCE OF BOLTZMANN AND GIBBS ENTROPY

According to Boltzmann, the system entropy in state {*N*_*A*_,*N*_*B*_} is S = k_B_·ln(Ω_NA,NB_), where Ω_NA,NB_ is the number of possible ways that the chemical state {*N*_*A*_,*N*_*B*_} can be occupied (i.e., the number of microstates). Specifically, Ω_NA,NB_ = *N*!/(*N*_*A*_!*N*_*B*_!). The change in entropy with a single chemical step from {*N*_*A*_,*N*_*B*_} to {*N*_*A*_–1,*N*_*B*_+1} is for a forward step,

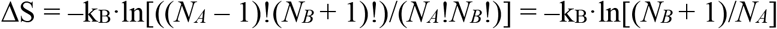

and for a reverse step

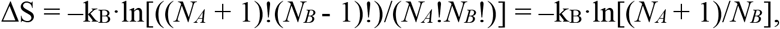

where the net change in entropy for a forward step from {*N*_*A*_,*N*_*B*_} to {*N*_*A*_–1,*N*_*B*_+1} followed by a reverse step from{*N*_*A*_–1,*N*_*B*_+1} to {*N*_*A*_,*N*_*B*_} is zero. The change in entropy with larger chemical steps (δη > 1) including molar steps is considered in the Appendix.

Because *N* individual switches (e.g., isolated in separate single molecule experiments) do not constitute a binary system,

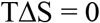

Thus, with thermal scaling from *N* individual molecules to an ensemble of *N* molecules, ensemble entropy is created, where

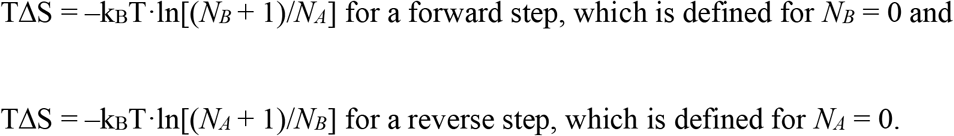

## VII. REACTION FREE ENERGY EQUATIONS

According to Eqs. 1 and 2, the reaction free energy for *N* individual molecules is

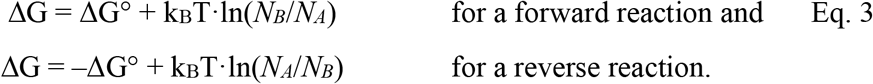

and the reaction free energy for an ensemble of *N* molecules is

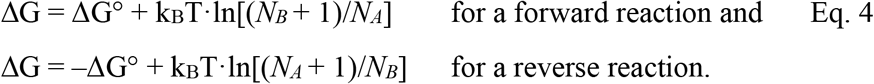

Thus, thermal scaling from the former to the latter physically creates a statistical state occupancy, k_B_T·ln[(*N*_*B*_ + 1)/*N*_*A*_], with a loss of a physical state occupancy, k_B_T·ln(*N*_*B*_/*N*_*A*_), and it does so smoothly (Eqs. 3 and 4 are incrementally different).

On the thermal scale of a single molecule the free energy change for a single chemical step is

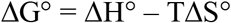

And on the thermal scale of an ensemble of *N* molecules the free energy change for a single chemical step is

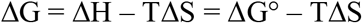

A comparison of these two equations shows that thermal scaling physically transforms ΔH° – TΔS° on the scale of a molecule into ΔH on the scale of an ensemble of molecules. Thus, entropy contributes to a measurable ΔH on a thermal scale on which that entropy is not defined. This is precisely why, when one fails to understand that thermal scaling is physical, entropy is perceived as a spook and described as “missing information” [15,24,25].

Here I describe how thermal scaling physically transforms the chemical kinetics of *N* individual switches into the chemical kinetics of an ensemble of *N* molecules described by thermal fluctuations within the ensemble energy landscape.

## VIII. THE ENSEMBLE REACTION ENERGY LANDSCAPE

Figure 2B illustrates entropic changes along the reaction coordinate for an ensemble of *N* = 5 molecules. When the system is in state {5,0}, Ω_5,0_ = 1. Unlike molecular models where k_B_·ln(0/5) is undefined, here the system entropy, k_B_·ln(1), is zero. After one net forward step, the system enters state {4,1}, increasing the number of microstates from 1 to 5 and increasing the system entropy from 0 to k_B_·ln(5). This increase in the number of microstates physically pulls the reaction forward since there are five-times more micropathways in the forward direction than in the reverse direction. This contrasts with the chemical activity of molecules pushing a reaction forward. When ΔG^o^ = 0, the reaction continues until a maximum system entropy is reached along the reaction coordinate, equilibrating in system state {3,2} when (*N*_*B*_ + 1)/*N*_*A*_ = exp(0) (Eq. 4).

Figure 3A is a plot of the reaction free energy (Eq. 4) along the system reaction coordinate,{*N*_*A*_,*N*_*B*_}, for *N* = 10 molecules. The reaction entropy, k_B_T·ln[(*N*_*B*_+1)/*N*_*A*_], increases logarithmically with each step whereas the molecular free energy, ΔG^o^, is a constant offset (it is the same for each step). Figure 3A shows that when ΔG^o^ decreases from 0 to –1.5 k_B_T (down arrow) the equilibrium state of the system changes from {5.5,4.5} to {2,8} (right arrow) corresponding to a change from (*N*_*B*_ + 1)/*N*_*A*_ = exp(0) to (*N*_*B*_ + 1)/*N*_*A*_ = exp(1.5). Because ΔG is the change in free energy, G, of the system with a chemical step (Eqs. 1 and 2), the free energy, G, of the system can be obtained by integrating Eq. 2 over the reaction coordinate. In Fig. 3B, the two plots in Fig. 3A are integrated and replotted as free energy, G. These are the system free energy landscapes described by Gibbs within which a reaction equilibrates at the point along the reaction coordinate{*N*_*A*_,*N*_*B*_} where G is a minimum. Figure 3 clearly shows that entropic forces, TΔS, (Fig. 3A) drive a reaction down an entropic energy funnel, TS (Fig. 3B) toward equilibrium.

**Fig. 3.**
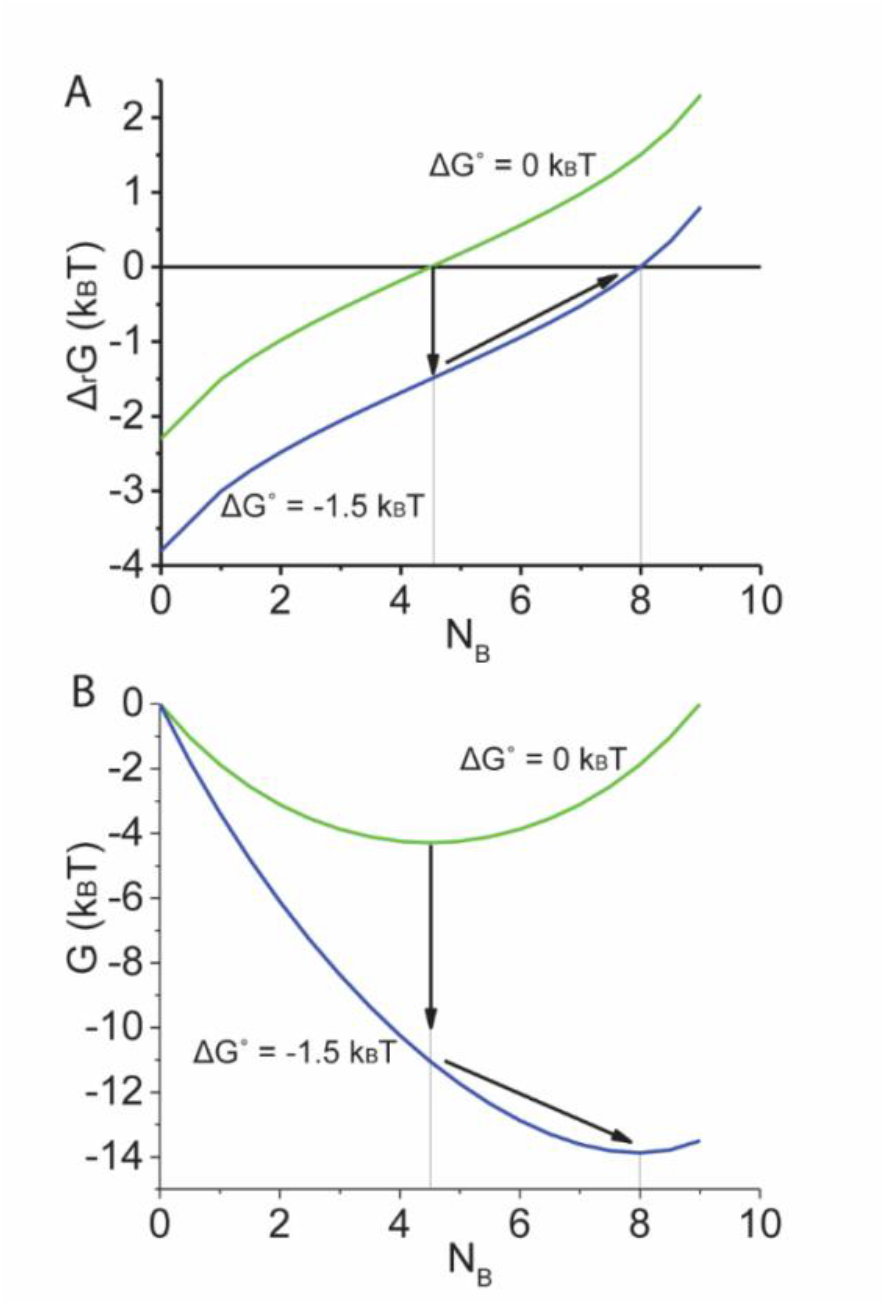
Change in Free energy along a reaction coordinate. **(A)** Equation 5 (*N* = 10 and *F* = 0) is plotted for a ΔG° of 0 (green line) and –1.5 k_B_T (blue line). For ΔG° = 0, the reaction equilibrates (Δ_r_G = 0) at {5.5,4.5} (x-axis = 4.5, y-axis = 0). When ΔG° decreases from 0 to –1.5 k_B_T (down arrow) the system re-equilibrates (right arrow) at {2,8} (x-axis = 8, y-axis = 0). **(B)** The system free energy, G, (integrals of Δ_r_G in panel A over the reaction coordinate) is plotted for ΔG° of 0 (green line) and –1.5 k_B_T (blue line). The same trajectory in panel A describing re-equilibration following a decrease in ΔG° is illustrated in panel B (arrows).

Both equilibrium and non-equilibrium kinetics and energetics are described by diffusion within the molecular energy landscapes of arbitrary molecules (ΔG^o^, Fig. 1B) tilted by an ensemble entropic energy landscape (TΔS, Fig. 3B). Because *N*_*A*_ and *N*_*B*_ are reaction coordinates, they do not affect the entropic landscape, which is to say the physical presence of molecules (*N*_*A*_ and *N*_*B*_) has no effect on the kinetics of a chemical reaction. Instead, kinetics are entropically driven. Like free energy (Eqs. 3 and 4), chemical kinetics are mathematically similar for both *N* individual molecules and an ensemble of *N* molecules (see Appendix) even though they describe physically different mechanisms. This analysis is presented in the Appendix and illustrated in Fig. 4.

**Fig. 4.**
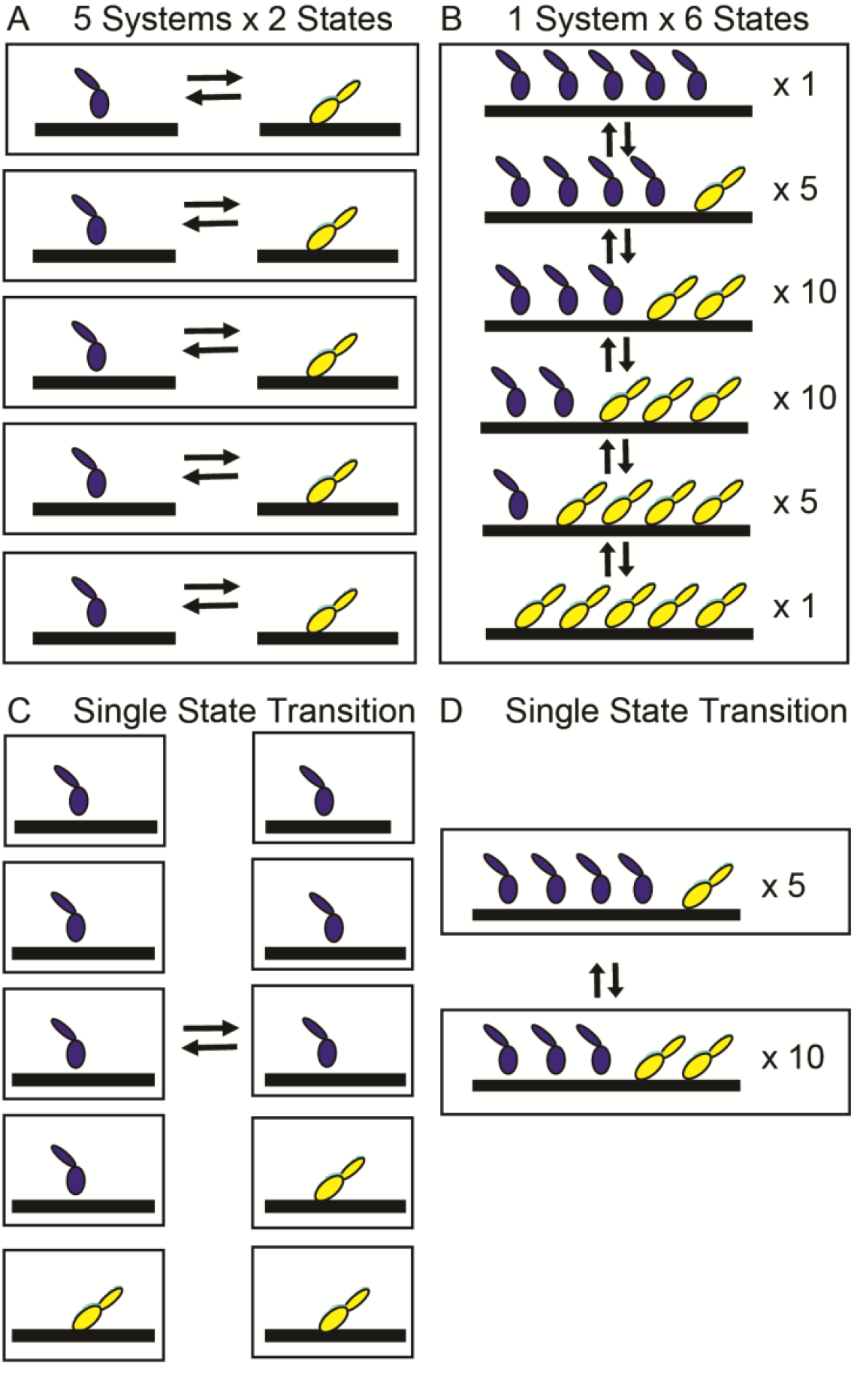
Five independent molecular switches versus an ensemble of five molecular switches. Thermal fluctuations isolated within five independent molecular switches (Figs. 1A and 2A) are described as five thermodynamic systems each with two states. **(B)** Thermal fluctuations among five molecular switches (Figs. 1A and 3B, only with five molecules) are described as one ensemble thermodynamic system having six states within which there are 1, 5, or 10 microstates (right). (C) A single state transition for five independent molecular switches. (D) A single state transition for an ensemble of five molecular switches.

## IX. HOW KINETICS AND ENERGETICS SCALE FROM *N* INDIVIDUAL MOLECULES TO AN ENSEMBLE OF *N* MOLECULES

Figure 4 compares the kinetics of 5 independent molecular switches (Fig. 4A) and the kinetics of an ensemble of 5 molecular switches (Fig. 4B). In Fig. 4A, each molecular switch is its own independent thermodynamic system each having two states. Figure 4B is a single ensemble system consisting of five molecular switches with six states (Fig. 3B) where each state has a different number of possible ways it can be occupied (Ω_NA,NB_ of 1, 5 or 10 microstates).

Figure 4C illustrates a molecular state transition from A to B of one out of the five independent molecular switches (from *N*_*A*_ = 4; *N*_*B*_ = 1 to *N*_*A*_ = 3; *N*_*B*_ = 2). The forward and reverse rates for this reaction, 4*k*°_*+*_ and 2*k*°_–_, depend on the physical numbers of molecules in states A and B. The ratio of these rates is (4/2)·(*k*°_*+*_/*k*°_–_), and the non-equilibrium reaction free energy is –k_B_T·ln[(4/2)·(*k*°_*+*_/*k*°_–_)] = ΔG° – k_B_T·ln[(4/2)] = ΔG° – k_B_T·ln(*N*_*A*_/*N*_*B*_), consistent with the molecular analysis above.

Figure 4D illustrates an ensemble state transition from {4,1} to {3,2}. The forward and reverse rates for this reaction do not depend on *N*_*A*_ or *N*_*B*_, and according to Eq. S1 the ratio of these rates is (10/5)·(*k*°_*+*_/*k*°_–_). The non-equilibrium free energy equation is –k_B_T·ln[(10/5)·(*k*°_*+*_/*k*°_–_)] = ΔG° – k_B_T·ln(Ω_NA–1,NB+1_/Ω_NA,NB_) = ΔG° – TΔS, consistent with an entropic analysis.

While molecular (chemical activity) and ensemble (entropic) kinetics and energetics are mathematically similar, they describe fundamentally different systems. In a chemical kinetic analysis, *N*_*A*_ = 4 individual molecules push the reaction forward and *N*_*B*_ = 2 individual molecules push the reaction backward, and the probability that the physical presence of molecules push the reaction forward relative to pushing the reaction backward is 4/2 = 2. In an entropic analysis the number of microstates, Ω_NA,NB_ = 10, in state {3,2} pulls the reaction forward toward 10 options and the number of microstates, Ω_NA,NB_ = 5, in state {4,1} pulls the reaction backward toward 5 options, where the probability that entropy pulls the reaction forward relative to pulling the reaction backward is 10/5 = 2.

While mathematically equivalent, there is a clear physical difference between the kinetics of the flip of one of five switches in Figs. 4A and the flip of one of five switches in 4B. The former describes the physical occupancy of the states of molecular switches, *N*_*A*_ and *N*_*B*_, pushing the reaction forward (Fig. 4A), whereas the latter describes the number of ways, Ω_NA,NB_, an ensemble of switches can be flipped irreversibly pulling the reaction forward. Remarkably, the latter describes the arrow of time pulling the system into the future down a priori defined entropic funnels. While this may sound like speculative theoretical physics, it is in fact the experimentally observed chemical thermodynamic mechanism of muscle contraction.

## X. DISCUSSION

In a companion article, I describe the convergence of top-down (thermodynamic) physics and bottom-up (molecular) biology through thermal scaling. I show that when *N* individual molecules thermally scale up to an ensemble of *N* molecules, ensemble entropy is created within a resonant ensemble structure. Shown here, within a thermal scale, the irreversible kinetics of a change in chemical state of the ensemble occurs with an increase in the ensemble entropy.

I show that like chemical energetics, chemical kinetics transforms smoothly across thermal scales. The chemical states of *N* individual molecules, *N*_*A*_ and *N*_*B*_, are defined by the physical occupancy of those states. The occupancy of *N*_*A*_ and *N*_*B*_ contributes to free energy as the chemical activity of states A and B. A change in chemical state is described by a change in *N*_*A*_ and *N*_*B*_. The kinetics of this change is described by *N*_*A*_ and *N*_*B*_ physically pushing the reaction forward and backward (i.e., mass action). In contrast, the chemical states of an ensemble of molecules, {*N*_*A*_,*N*_*B*_}, is described by an a priori probability density function defined independent of the physical occupancy of states A and B. The a priori ensemble entropy of state {*N*_*A*_,*N*_*B*_} contributes to free energy. A change in chemical state is described by a change in {*N*_*A*_,*N*_*B*_} and a change in ensemble entropy. The kinetics of this change is described by the ensemble being physically pulled toward higher entropy.

These are not two alternative hypotheses. They are two physically different systems. The former physically collapses into the latter through thermal scaling with the creation of ensemble entropy. Within each thermal scale, the ensemble entropy created by components at a lower thermal scale contributes to the energy landscape of the ensemble system. For example, the kinetics and energetics of an individual protein are described by constants (ΔG°, ΔH°, ΔS°, *k*°_*+*_ and *k*°_–_) that are defined a priori in terms of its reaction energy landscape (Fig. 2A) independent the occupancy of the states of constituent structural parts. Likewise, the kinetics and energetics of a protein ensemble are described by constants (ΔG, ΔH, ΔS, *k*_*+*_ and *k*_–_) that are defined a priori in terms of its reaction energy landscape (Fig. 3B) independent of the occupancy of the states of its constituent protein parts. With the transformation of *N* molecular landscapes (Fig. 2A) into an ensemble landscape with *N* states (Fig. 3B) molecular wells are no longer defined for individual molecules and are physically subsumed by the entropic well of the ensemble.

This analysis solves several important paradoxes in physics. It describes observational uncertainty as originating when the statistical occupancy of states physically replaces the physical occupancy of molecular states when entropy is created with scaling. Thus, the mysteriousness of “not knowing” the occupancy of states on smaller scales embodied by both Schrodinger’s cat (on the energy scale of light) and Maxwell’s demon (on the energy scale of heat) is not mysterious at all; the perceived mysteriousness comes from our “not knowing” that scaling creates entropy through a physical and irreversible transformation. Thermal scaling describes entanglement as two physically connected systems that scale up in parallel with overlapping energy spacings (see companion article). It solves the Gibbs paradox [26]. It describes the arrow of time as the pull of a priori defined possible future states (entropy) rather than a push by states physically occupied in the present. It disproves all classical mechanical mechanisms of biological function. And it implies that life on earth evolved not in local opposition to the second law of thermodynamics but because of it.

Muscle evolved constrained by the physical laws of the universe – not physical laws as we understand them today – and so by dissecting muscle down to its constituent parts and studying how these parts come together across thermal scales we learn about universal physical laws. A.V. Hill showed in 1938 that the mechanics and energetics of muscle contraction are not defined by the mechanics and energetics of the molecules within muscle. This extraordinary result has been widely ignored because it is counter to the bottom-up corpuscularian philosophy of most molecular biologists. We have since shown experimentally that the physical basis for Hill’s observation is the entropy created in muscle when *N* individual molecular switches thermally scale up to an ensemble of *N* switches. The resulting two-state thermodynamic model of irreversible entropically-driven muscle power output accounts for most key aspects of muscle contraction [11,18,19]. In summary, we directly observe in muscle that thermal scaling is physical and that the entropy it creates is measurable, useful, and irreversibly pulls us into the future. While controversial, this is less fantastical than other widely varied mysterious depictions of scaling, entropy, and the arrow of time.

## XI. APPENDIX

### 1. Change in system entropy for δη > 1

The change in system entropy associated with δη chemical steps from {*N*_*A*_,*N*_*B*_} to {*N*_*A*_– δη,*N*_*B*_+δη} is

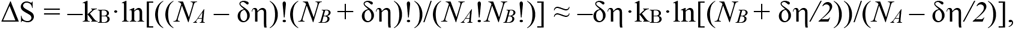

which defines an average change in system entropy, k_B_T·ln[(*N*_*B*_ + δη/2)/(*N*_*A*_ – δη/2)], over δη steps. The change in entropy, ΔS, relative to S = 0 occurs from {*N*/2,*N*/2} to {*N*/2–δη, *N*/2+δη}, or

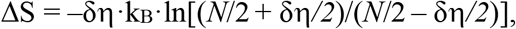

or

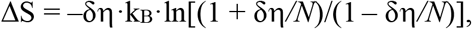

where δη*/N* is a fractional change in the extent of the reaction relative to unity. Assuming δη = 1 mol, this equation can be written in its molar form

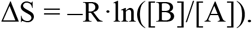

While this resembles the chemical activity difference between states and B defined in terms of physical quantities of molecules in states A and B, ΔS describes an a priori change in entropy defined independent of physical quantities of molecules.

### 2. Equilibrium Energetics

At equilibrium (Δ_r_G = 0), the probability of finding the system in state {*N*_*A*_+1,*N*_*B*_–1} relative to {*N*_*A*_,*N*_*B*_} is exp[–ΔG°/k_B_T + ln(Ω_NA-1,NB+1_/Ω_NA,NB_)], where ln(Ω_NA-1,NB+1_/Ω_NA,NB_) = TΔS/k_B_T. When ΔG° = 0, Ω_NA,NB_ is the probability density function, *P*_*NA*,*NB*_. The equilibrium constant, *K* = (Ω_NA,NB_/Ω_NA-1,NB+1_)_eq_ = exp[–ΔG°/k_B_T] is *K* = (*N*_*B(eq)*_ + 1)/*N*_*A(eq)*_ and describes the point along the reaction coordinate {*N*_*A*_,*N*_*B*_}_eq_ at which the system equilibrates (Fig. 3). For comparison, in chemical activity models the equilibrium constant *K* = *N*_*B(eq)*_/*N*_*A(eq)*_ describes the physical distribution of *N* molecules within the energy landscape of a single molecule (Fig. 2A).

### 3. Non-Equilibrium Energetics

When the system is perturbed from equilibrium by a change in the internal energy of the system, δE, [i.e., a change in ΔG° or k_B_T·ln[(Ω_NA-1,NB+1_/Ω_NA,NB_)], the free energy equation becomes

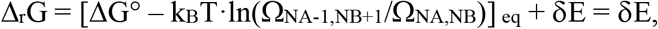

where δE is a non-equilibrium (ne) perturbation to any of the energy term on the right-hand side of Eq. 5. Here,

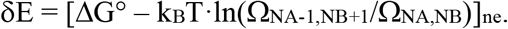

As illustrated in Fig. 3, if the system is perturbed by a change in ΔG°, the system relaxes to a new equilibrium state, {*N*_*A*_,*N*_*B*_}_eq_, as k_B_T·ln(Ω_NA-1,NB+1_/Ω_NA,NB_) approaches the new ΔG°.

Through this process, Δ_r_G returns to zero and δE remains in the system in the form of a change in system entropy. If the system perturbation is a change in k_B_T·ln(Ω_NA-1,NB+1_/Ω_NA,NB_) [an irreversible transfer of molecules between molecular states] the system relaxes back to the original equilibrium state, {N_A_,N_B_}_eq_, as k_B_T·ln(Ω_NA-1,NB+1_/Ω_NA,NB_)_ne_ approaches ΔG°. Through this process the entropic δE is lost from the system as heat. This analysis becomes more complex when non-PΔV work is performed, in which case, in addition to exchanges between k_B_T·ln(Ω_NA-1,NB+1_/Ω_NA,NB_) and ΔG°, δE can be exchanged with internal mechanical potentials; and in addition to being lost from the system as heat or stored in the system as entropy, δE can be lost from the system as work performed on the surroundings [11].

### 4. Equilibrium Kinetics

Chemical kinetics, like energetics, have both molecular and system components. The molecular component is the molecular kinetic rate constants, *k°*_*–*_ and *k°*_*+*_, defined a priori (independent of the occupancy of the landscape) by the molecular energy landscape in Fig. 2A. Here I show that the system kinetic component is the number of microstates into which the system is pulled, defined a priori (independent of the occupancy of the landscape) by the system energy landscape in Figs. 2B and 3. This contrasts with a chemical activity analysis in which the system kinetic component is a physical number of molecules that occupy and push molecules out of a state in a molecular landscape. I show that the two approaches are mathematically similar, but only the former entropic analysis allows one to simulate non-equilibrium kinetics in a thermodynamic system.

In general, the net rate at which the system transitions from chemical state {*N*_*A*_,*N*_*B*_} to chemical state {*N*_*A*_–1,*N*_*B*_+1} is

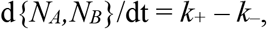

where *k*_*+*_ and *k*_–_ are the forward and reverse transition rates between these states. Again, this is not the rate at which molecules transition between states with units of number of molecules per second, it is the rate at which the system transitions between ensemble chemical states along the reaction coordinate {*N*_*A*_,*N*_*B*_} of the system energy landscape (Figs. 2B and 3) with units of chemical states per second. If *k*_*+*_ > *k*_–_, the reaction progresses left to right along the reaction coordinate of system energy landscape at a positive rate *k*_*+*_ – *k*_–_. If *k*_*+*_ < *k*_–_, the reaction progresses right to left along the reaction coordinate of the system energy landscape at a positive rate *k*_–_ – *k*_*+*_. If *k*_*+*_ = *k*_–_, the reaction is at equilibrium.

At equilibrium, the probability of finding the system in state {*N*_*A*_+1,*N*_*B*_–1} relative to state {*N*_*A*_,*N*_*B*_} is

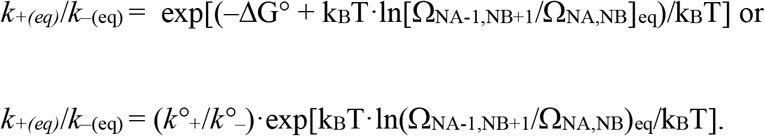

Here, *k°*_*+*_ and *k°*_*–*_ are molecular contributions to *k*_*+*_ and *k*_–_ and exp[–TΔS/k_B_T] is the system contribution to *k*_*+*_ and *k*_–_. The influence of –ΔS/k_B_ on kinetics can be understood energetically as a tilt of the system landscape that adds to the tilt of the molecular landscape, ΔG°. It can also be understood kinetically as

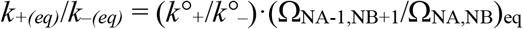

where the factor Ω_NA-1,NB+1_/Ω_NA,NB_ describes the number of micropathways available for the forward reaction relative to the reverse reaction. For example, in Fig. 2B there are five micropathways from {5,0} to {4,1} and one micropathway back from {4,1} to {5,0}, which means that the forward reaction is entropically five-fold more likely than the reverse reaction (Ω_NA-1,NB+1_/Ω_NA,NB_ = 5). This implies forward and reverse rate constants of

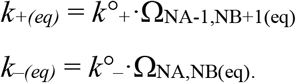

At equilibrium, *k*_*+(eq)*_ = *k*_*–(eq)*_, or

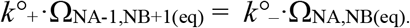

Recall that

ΔS = –k_B_·ln(Ω_NA,NB_/Ω_NA-1,NB+1_) = –k_B_·ln [(*N*_*B(eq)*_ + 1)/ *N*_*A(eq)*_) for a forward step and ΔS = –k_B_·ln(Ω_NA-1,NB+1_/Ω_NA,NB_) = –k_B_·ln [(*N*_*A(eq)*_ + 1)/ *N*_*B(eq)*_) for a reverse step and on comparison,

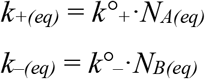

which is mathematically consistent with a chemical activity analysis, only here again *N*_*A(eq)*_ and *N*_*B(eq)*_ are not physical numbers of molecules but the system reaction coordinate {*N*_*A*_,*N*_*B*_}_eq_ at which the system equilibrates.

### 5. Non-Equilibrium Kinetics

Chemical kinetics is fully determined from the molecular and system tilt of the system energy landscape. It follows that non-equilibrium kinetics is fully determined by the non-equilibrium tilt, δE, of these landscapes, where a non-equilibrium perturbation to ΔG° affects the molecular rates *k°*_*+*_/*k°*_*–*_, and an irreversible transfer of molecules between molecular states affects Ω_NA,NB_/Ω_NA-1,NB+1_. In general,

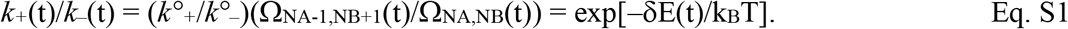

According to Eq. 10, when δE(t) < 0, *k*_*+*_(t) > *k*_*–*_(t), consistent with the requirement that a negative δE(t) drives the reaction forward. This forward reaction increases TΔS (Fig. 3) until δE(t) = 0, at which point *k*_*+*_(t) = *k*_*–*_(t). When δE(t) > 0, *k*_*+*_(t) < *k*_*–*_(t), consistent with the requirement that a positive δE(t) drives the reaction backward. This reaction decreases TΔS (Fig. 3) until δE(t) = 0, at which point *k*_*+*_(t) = *k*_*–*_(t).

The non-equilibrium rate equation is the same as the equilibrium rate equation

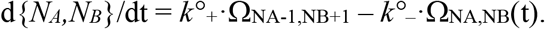

For comparison with a chemical activity analysis, from above

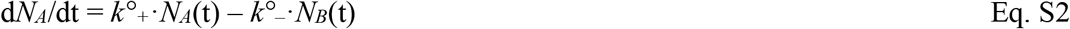

where *N*_*A*_(t) and *N*_*B*_(t) describe the time dependence of steps along the system reaction coordinate {*N*_*A*_,*N*_*B*_}, not the time-dependence of the physical numbers of molecules in different states. Changes in *N*_*B*_(t) + 1 are equal and opposite changes in *N*_*A*_(t) along the system reaction coordinate, and so

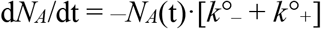

which means that *N*_*A*_(t) decreases exponentially with time as

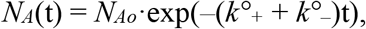

with a relaxation rate, *k*°_*+*_ + *k*°_–_, where *N*_*A*o_ is the initial position on the reaction coordinate {*N*_*A*_,*N*_*B*_}_o_.

## Acknowledgments

I thank LEB, JWG, Julie, and my students, colleagues, and mentors who have over many years inspired and guided this work. This work was funded by a grant from the National Institutes of Health 1R01HL090938-01.

## Data and materials availability

All data are available in the main text.

## REFERENCES

[1] S. Carnot, Reflections on the Motive Power of Fire (Dover, New York, 2005).

[2] A. V. Hill, The Heat of Shortening and the Dynamic Constants of Muscle, Proc. R. Soc. London. Ser. B 126, 136 (1938).

[3] S. Walcott, S. Sun, E. Debold, and W. Herzog, In Defense of Huxley, Biophys. J. 123, 1 (2024).

[4] J. Langer, Statistical Theory of the Decay of Metastable States, Ann Phys 54, 258 (1969).

[5] S. Chandrasekhar, Stochastic Problems in Physics and Astronomy, Rev Mod Phys 15, 1 (1943).

[6] H. Kramers, Brownian Motion in a Field of Force and the Diffusion Model of Chemical Reactions, Physica 7, 284 (1940).

[7] J. E. Baker, I. Brust-Mascher, S. Ramachandran, L. E. LaConte, and D. D. Thomas, A Large and Distinct Rotation of the Myosin Light Chain Domain Occurs upon Muscle Contraction., Proc. Natl. Acad. Sci. U. S. A. 95, 2944 (1998).

[8] J. E. Baker, L. E. W. LaConte, I. Brust-Mascher, and D. D. Thomas, Mechanochemical Coupling in Spin-Labeled, Active, Isometric Muscle., Biophys. J. 77, 2657 (1999).

[9] J. E. Baker and D. D. Thomas, A Thermodynamic Muscle Model and a Chemical Basis for A.V. Hill’s Muscle Equation, J Muscle Res Cell Motil. 21, 335 (2000).

[10] J. E. Baker, Muscle Contracts with the Flip of an Ensemble of Switches, Biophys. J. 74, A22 (1998).

[11] J. E. Baker, Thermodynamics and Kinetics of a Binary Mechanical System: Mechanisms of Muscle Contraction, Langmuir 38, 15905 (2022).

[12] C. Kittel and H. Kroemer, Thermal Physics (1980).

[13] A. Humeau-Heurtier, Multiscale Entropy Approaches and Their Applications, Entropy 22, (2020).

[14] F. Herrmann and M. Pohlig, Which Physical Quantity Deserves the Name “Quantity of Heat”?, Entropy 23, (2021).

[15] H. U. Fuchs, M. D’Anna, and F. Corni, Entropy and the Experience of Heat, Entropy 24, (2022).

[16] O. Darrigol, The Gibbs Paradox: Early History and Solutions, Entropy 20, (2018).

[17] A. Wehrl, General Properties of Entropy, Rev. Mod. Phys. 50, 221 (1978).

[18] J. E. Baker, Four Phases of a Force Transient Emerge From a Binary Mechanical System, J. Muscle Res. Cell Motil. (2024).

[19] V. Murthy and J. Baker, Stochastic Force Generation in an Isometric Binary Mechanical System, BioRxiv (2023).

[20] J. E. Baker, The Problem with Inventing Molecular Mechanisms to Fit Thermodynamic Equations of Muscle, Int. J. Mol. Sci. 24, (2023).

[21] H. Frauenfelder, S. G. Sligar, P. G. Wolynes, H. Frauenfelder, S. G. Sligar, and P. G. Wolynes, The Energy Landscapes and Motions of Proteins, 254, 1598 (1991).

[22] J. Howard, Mechanics of Motor Proteins and the Cytoskeleton, 1st ed. (Sinauer Associates, 2001).

[23] L. Stryer, Biochemistry (W.H. Freeman and Company, New York, 1995).

[24] C. E. Shannon, A Mathematical Theory of Communication, Bell Syst. Tech. J. 27, 379 (1948).

[25] R. H. Swendsen, Probability, Entropy, and Gibbs’ Paradox(Es), Entropy 20, (2018).

[26] J. E. Baker, Cells Solved the Gibbs Paradox by Learning to Contain Entropic Forces, Sci. Rep. 13, (2023).

